# Structures of co-transcriptional RNA capping enzymes on paused transcription complex

**DOI:** 10.1101/2023.08.09.552658

**Authors:** Yan Li, Qianmin Wang, Yanhui Xu, Ze Li

## Abstract

Nascent pre-mRNA undergoes 5′ end capping as the first step of processing. Early evidences demonstrated the guanosine addition and 2′-O-ribose methylation spatiotemporally correlated with transcription machinery at the early stage of transcription. Here, we determined cryo-EM structures of PEC (paused elongation complex)-RNGTT (RNA guanylyltransferase and 5′ phosphatase) and PEC-RNGTT-CMTR1 (cap-specific mRNA (nucleoside-2′-O-)-methyltransferase). The structures show that RNGTT docks to the root of Pol II stalk through its OB fold. Within RNGTT, the OB fold binds N-terminal of triphosphatase domain and facilitates positioning its catalytic cavity facing towards the RNA exit tunnel. RNGTT dephosphorylates and guanylates PEC-bound RNAs of 17nt, 19nt, 20nt, but not 22nt, in length. CMTR1 arrayed with RNGTT on the Pol II surface through distinct interfaces. Our structures unravel that capping enzymes RNGTT and CMTR1 directly docks to paused elongation complex, and shed light on how pre-mRNA capping couples with Pol II at the specific transcription stage.

## Introduction

Transcription is one of the most important biological processes, by which genetic information is transcribed from DNA to RNA and translated into protein for specific physiological functions^1,2^. Precursor messenger RNA (pre-mRNA), miRNA, siRNA and other non-coding RNAs are transcribed by RNA polymerase II^3^. Co-and post-transcriptional modifications including 5′ capping, splicing and 3′ polyadenylation maturate pre-mRNA into mRNA and distinguish it from other RNA products.

In eukaryotes, the unique 5′ cap structure, m^7^GpppNm, protects nascent pre-mRNA from degradation by ribonucleases, DXO^4^ and XRN^5^, and also facilitates mRNA nuclear export and translation^6,7^. In metazoans, m^7^GpppN, named Cap0, is stepwise synthesized by RNA guanylyltransferase and 5′-phosphatase (RNGTT) and RNA guanine-7 methyltransferase (RNMT) bound with its activator RAM^8-10^. RNGTT possesses a 5′-triphosphatase domain (TPase) and an RNA guanylyltransferase domain (GTase). TPase primarily removes the γ-phosphate of 5′-triphosphate and GTase adds the GMP to the 5′-diphosphate, forming the 5′ GpppN structure. Subsequently, RNMT-RAM methylates the N7 position of guanine to form the Cap0 structure ^11^. Specific RNA 2′-O ribosylmethyltransferases cap methyltransferase 1 (CMTR1) and 2 (CMTR2) methylate 2′-O of +1 and +2 ribonucleotides into m^7^GpppNm (Cap1) and m^7^GpppNmNm (Cap2) structures, respectively^12-14^.

The dephosphorylation, guanylation, and methylation do not occur until the nascent pre-mRNA protrudes from RNA exit tunnel. RNase footprinting assay showed that Pol II protects pre-mRNAs of 15 nucleotide (nt), which has been virtualized by the structures of transcription elongation complex (EC) ^15-17^. It is confirmed that 5′ end of the short transcripts is inaccessible for the capping enzymes. Capping begins at the 5′-triphosphate of the first transcribed nucleotide stretching out of Pol II, but 16 nt RNA is not qualified ^18-20^. It is demonstrated that capping does not occur until the transcript is elongated to 20 nt *in vitro* and 20-30 nt *in vivo*^21,22^

Previous studies have suggested that Pol II C-terminal domain (CTD) is involved in recruiting and activating the capping enzymes ^23,24^. The conserved Pol II CTD consists of a tandemly repeated consensus sequence Y^1^S^2^P^3^T^4^S^5^P^6^S^7^, which is subject to extensive phosphorylation. Phosphorylated Pol II CTD directly binds TPase and GTase ^25,26^.The Ser-5 phosphorylation catalyzed by CDK7 dominates early transcription stage in favor of recruiting and stimulating the guanylation activity of GTase ^27^. The presence of Ser-5 phosphorylated Pol II modestly increases the GMP-capping enzyme intermediate formation. Moreover, the CTD-independent capping activity has also been detected *in vitro* ^28^.

The DSIF (DRB sensitivity-inducing factor) component, SPT5, binds full-length RNGTT *in vitro*, in which TPase and GTase separately bind SPT5 ^26,29,30^. SPT5 stimulates guanylation activity of GTase to several folds with no effects on the 5′-triphosphatase activity ^31^. Interaction between SPT5 with the TPase domain has an allosteric effect on guanylation activity of GTase. Although phosphorylated CTD and SPT5 individually stimulates capping activity, their positive effects do not collaboratively intensify^32^. Besides the capping enzyme, SPT5 also recruits the negative elongation factor (NELF) to Pol II assembling into paused elongation complex (PEC) ^33^. DSIF and NELF repress elongation at the promoter-proximal region. In metazoans, transcribing Pol II stalled around 20-60 nt in nascent pre-mRNA^34,35^, where capping modification takes place. Capping occurs progressively from uncapped transcripts at early pausing state to more capped ones at late pausing state, illustrating that 5′ end capping and Pol II pausing are spatiotemporally coupled rather than isolated ^19^.

Cap0 and Cap1 modifications are accomplished in nucleus, whereas three enzymes vary in the genomic patterns. CMTR1 is enriched at 5′ end of genes with its peak proximal to the TSS, nevertheless, RNMT-RAM distributes along the entire pre-mRNA, which suggests CMTR1 possibly more connects with early stage transcription events than RNMT-RAM^10,36^. The C-terminal WW domain of CMTR1 interacts with the Ser-5 phosphorylated CTD ^37^, indicating that CMTR1 is closely associated with the transcribing Pol II and CMTR1 may simultaneously dock on the Pol II with RNGTT.

Although structures of poxvirus and yeast co-transcriptional capping enzymes have been determined, the components of mammalian co-transcriptional capping complex are proved to differ from others. In *Vaccinia* poxvirus, a single trifunctional protein (D1) possesses triphosphatase, guanylyltransferase and methyltransferase activities, forming a heterodimer with its activator D12 to produce the Cap0 structure in complex with vRNAP ^38^. In *S. cerevisiae*, the capping enzyme is composed of triphosphatase Cet1 and guanylyltransferase Ceg1, which form hetero-trimer or hetero-tetramer that binds transcribing Pol II ^39^. For metazoan capping enzyme, only crystal structures of isolated domains have been determined till now ^23,40^. Moreover, several transcriptional factors get involve in metazoan capping enzyme regulation ^41^. The functional architecture of full-length human capping enzyme, especially in the context of mammalian Pol II with other transcription factors remains elusive. Here, we employed *S. scrofa* Pol II, human DSIF, NELF, RNGTT, and CMTR1, and determined PEC-RNGTT and PEC-RNGTT-CMTR1 complex structures at nominal resolution 3.53 Å and 4.0 Å, respectively. In PEC-RNGTT, we unveil that RNGTT is docked adjacent to the Pol II stalk with the OB fold inserting into the root of stalk, and positions TPase at the RNA exit tunnel. In PEC-RNGTT-CMTR1, we represented the RNGTT and CMTR1 arrayed at the Pol II periphery in the aid of different interfaces of the Pol II stalk, neighboring the RNA exit tunnel.

## Results

### Assembly of the PEC-RNGTT complex

Referring to the temporal and spatial correlation of pre-mRNA capping with early stage of transcription, previous studies have shown that Ser-5 phosphorylated Pol II CTD ^23,24^ and DSIF^24,32^ stimulate capping enzymatic activity, and NELF-mediated Pol II pausing tightly relates with capping events^19^. We first tested whether RNGTT binds Pol II dependent of CTD phosphorylation. In a glycerol gradient centrifuge assay, the purified RNGTT protein co-migrated with Pol II that was subjected to CTD phosphorylation by human transcription factor TFIIH ^42^, and no co-migration was observed for the unphosphorylated Pol II (Supplementary Figs. 1a, d). We therefore used phosphorylated Pol II in complex assembly. Considering the stimulation of SPT5 to capping enzymatic activity, we next tested the role of DSIF in RNGTT binding with the phosphorylated Pol II in the absence and presence of NELF subcomplex, respectively. DSIF by itself or in combination with NELF did not disrupt RNGTT-Pol II interactions, and seemed to co-migrate with RNGTT-Pol II to fractions of higher molecular weight. The addition of NELF complex had no negative effect on RNGTT binding to Pol II (Supplementary Figs. 1b, e). We therefore assembled the complex by sequential addition of phosphorylated Pol II, RNA-DNA hybrid, non-template strand, DSIF, RNGTT and NELF.

To choose pre-mRNA in a proper length for complex assembly, we tested the capping enzymatic activity with RNGTT in the context of Pol II elongation complex. Using *in vitro* transcribed 17 nt, 19 nt, 20 nt, 22 nt 5′-triphosphate pre-mRNA, RNGTT efficiently guanylated the 19nt (53.3%) and 20nt (51.1%) 5′-triphosphate RNA substrates, and the 17 nt RNA (48.9%) showed comparable capping efficiency to 19 nt and 20 nt. However, 22 nt 5′-triphosphate pre-mRNA was hardly guanylated in our guanylation assay (Fig. 2g). According to the length of pre-mRNA accommodated in the RNA tunnel in high-resolution Pol II structures ^15-17,43^, we chose 17 nt 5′-triphosphate RNA as the nascent 5′-triphosphate pre-mRNA substrate, which was successfully assembled into the paused elongation complex (PEC) in the presence of RNGTT for cryo-EM data collection.

### Structure determination of the PEC-RNGTT complex

Cryo-EM structure of PEC-RNGTT complex was determined from 4,106,085 particles at a nominal resolution of 3.5 Å (Supplementary Figs. 2b, d). Cryo-EM 3D reconstructions showed that RNGTT locates adjacent to the Pol II stalk, and NELF complex around the Pol II funnel is discussed later. The Pol II core was resolved at 3 Å, whereas DSIF, NELF and RNGTT at periphery were solved at local resolution of 5-7 Å (Fig.1b and Supplementary Fig. 2d). Structural model of PEC (PDB ID:6GML) ^43^ was rigidly fitted into the corresponding densities, and manually adjusted in Coot (Supplementary Fig. 3a).

**Fig. 1.**
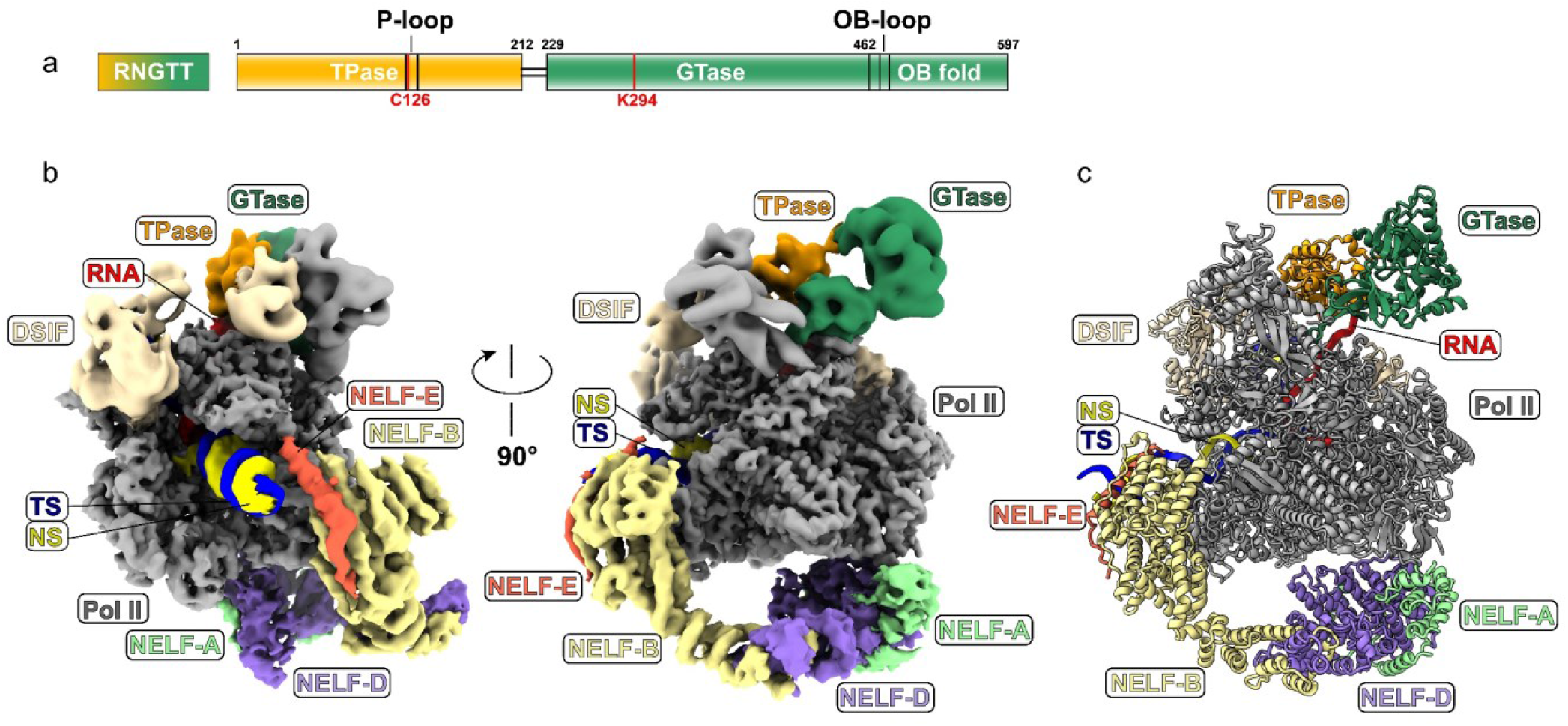
Overall cryo-EM structure of the PEC-RNGTT. **a** Domain architecture of RNGTT. The cryo-EM color code is used throughout all figures. Solid black lines represent the linkage region between TPase and GTase domains of RNGTT. The active site residues of the enzyme are marked in red lines. **b** Combined cryo-EM map of PEC-RNGTT in two different views and **c** Structural model of PEC-RNGTT in the same view. All components are depicted in corresponding tab color.

Focused refinement notably improved the resolution of RNGTT to 5.8 Å (Supplementary Fig. 2b). The moderate resolution of RNGTT is consistent with earlier structural studies showing that the capping enzymes are highly mobile on Pol II ^39^. Structural model of the GTase (PDB ID:3S24)^23^ was docked into the locally refined cryo-EM map, in which the density of the OB fold was clearly visualized (Fig. 1a and Supplementary Fig. 3b). The continuous density at two-domain boundary and the N-terminal direction of GTase gave us a clue to place TPase model into the map (Figure 1a and Supplementary Fig. 3c). We readily positioned the models of TPase of RNGTT (PDB ID:1I9S), GTase of RNGTT (PDB ID:3S24) and NELF moiety (PDB ID:6GML) into corresponding maps and adjusted the model in Coot. The 5′ end of 17 nt pre-mRNA was observed in the local refined map (Fig. 1b and Supplementary Fig. 3d), and we finally imaged a whole picture of PEC-RNGTT complex (Fig. 1c and Supplementary Fig. 3a)

### Structure of the PEC-RNGTT complex

The structure of PEC-RNGTT reveals PEC similar to the previously determined PEC structure^43^. DSIF domains wrap around the Pol II body from the upstream DNA entry to nascent RNA exit tunnel. NELF complex hung at the periphery of the Pol II body (Figs. 1b, c and Supplementary Fig. 3a). One copy of RNGTT is located adjacent to the stalk of Pol II (Figs. 1b and 2b). The OB fold, characterized by binding with oligonucleotide/oligosaccharide, mediates RNGTT-stalk interactions and stretches its OB loop (residue 476-484 aa) into the gap between RPB1 and RPB7 (Fig. 2c). The OB fold is well-positioned on the Pol II stalk and seems to bring RNGTT in close proximity to the RNA exit tunnel, rather than the previously proposed Pol II foot ^44^.

**Fig. 2.**
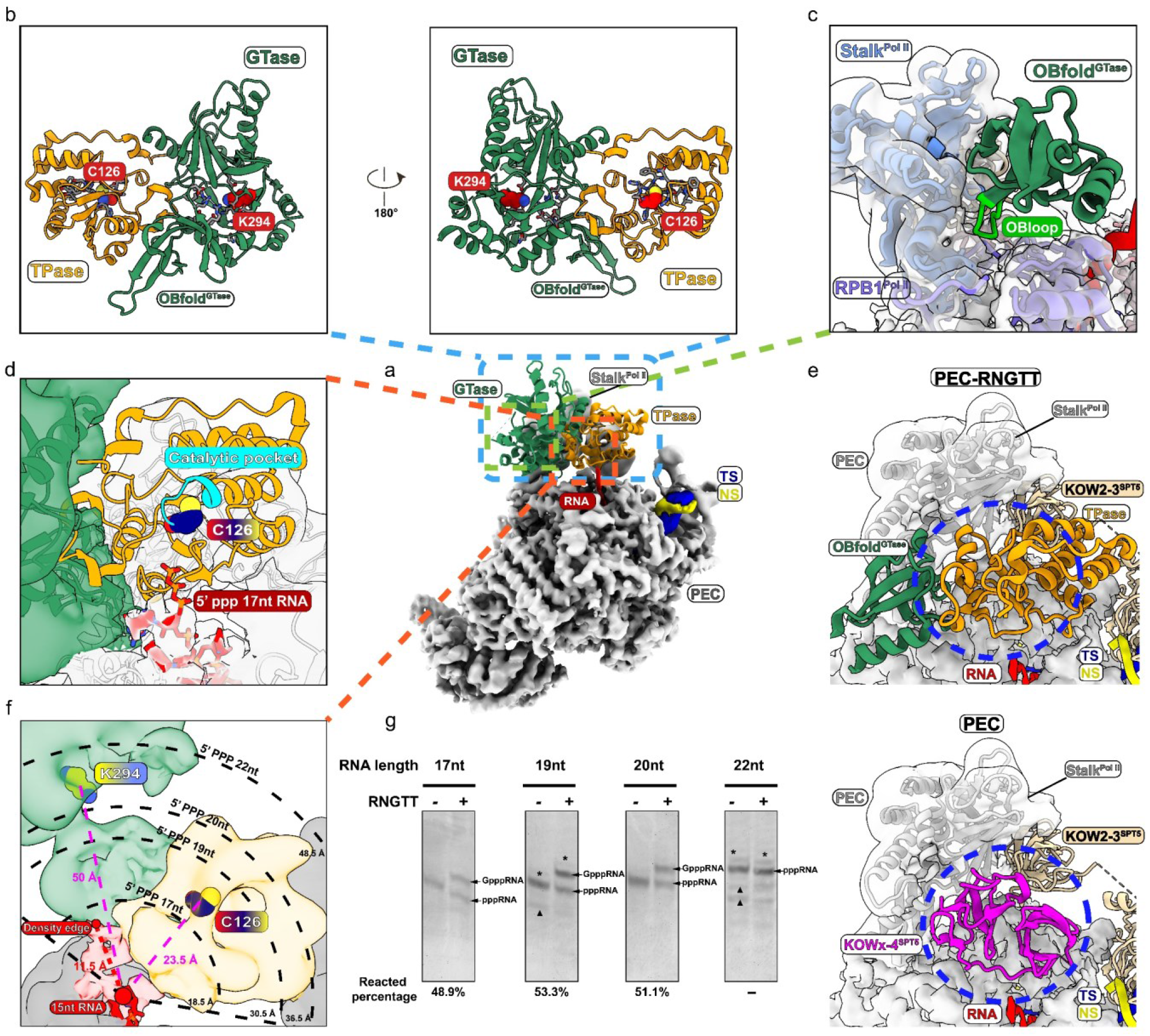
RNGTT docks in close proximity to the Pol II stalk and catalyzes 5′-triphosphate RNAs in different lengths. **a** Ribbon model of RNGTT docks on the PEC cryo-EM map. TPase and GTase domains of RNGTT are represented as ribbons in orange and sea green respectively while other components as density maps with labels in the corresponding tab colors. **b** Close-up view of of TPase and GTase domains of RNGTT. The active centers are displayed as spheres and colored by elements. **c** OB fold of GTase interacts with Pol II stalk (cornflower blue) and RPB1 subunit (medium slate blue). Pol II is illustrated as grey density map. OB loop of OB fold is highlighted in lime. **d** TPase domain lies at the RNA exit tunnel. The active center of TPase is displayed as spheres and colored by elements. The triphosphate of RNA is displayed as sticks. Oxygen is colored in red and phosphorus is in orange. **e** Comparison of PEC-RNGTT (above) and PEC (below) in the same view. (Pol II, grey; DSIF, wheat; KOWx-KOW4 domain, magenta). The blue ellipse circled clash position between KOWx-KOW4 and RNGTT. f Schematic diagram of distances between 5′ end of 15 nt RNA and two active sites of RNGTT. The 5′ end of 15 nt RNA is marked as the larger red circle and the end of visible RNA density edge is marked as the smaller red circle, which are linked by red dashes. The active residues of the enzymes are displayed as spheres and colored by elements. The dashes in magenta display distances between 15nt RNA end and enzymic active centers and the dotted circular sectors display estimated ranges of 5′-triphosphate RNAs in 17 nt, 19 nt, 20 nt and 22 nt. **g** *In vitro* guanylation assay of 17 nt, 19 nt, 20 nt and 22 nt 5′-triphosphate RNA in the context of phosphorylated Pol II-DSIF-RNGTT complex. Product percentages quantified by ImageJ are labeled below the lanes. There is no obvious band for 22 nt GpppRNA, therefore its product percentage is written as (-). Contamination bands are marked with asterisks and degradation bands with triangles.

In our cryo-EM structure, the TPase domain not only tiles with GTase domain, but also interacts with the OB fold through the N-terminal of TPase, forming a head-to-tail intra-molecular contact (Fig. 2b). TPase domain of RNGTT is surrounded by the OB fold, KOW2-3 of SPT5 and the Pol II stalk with the catalytic pocket facing towards the pre-mRNA exit tunnel (Figs. 2c, d and e).

Previous studies suggest that KOWx-KOW4 and KOW5 form an RNA clamp at RNA exit tunnel, which protects nascent pre-mRNA for subsequent processing^15,43^. In PEC-RNGTT structure, binding of RNGTT disrupts the original interface between KOWx-KOW4 and Pol II, and the position of KOWx-KOW4 in PEC is occupied by TPase domain and part of OB fold (Fig. 2e). We hypothesized that, in PEC-RNGTT, OB fold interplays with the stalk, resulting in the KOWx-KOW4 destabilization and the placement of TPase of RNGTT at RNA exit tunnel for 5′ end capping.

### PEC-RNGTT could guanylate 17 nt RNA substrate in vitro

The minimal length of RNA for capping reaction has been controversial for decades. In general, the minimum length for nascent pre-mRNA to be capped is around 20 nt, and longer products as 79 nt and 223 nt could be capped as efficiently as short transcripts^18,22,28^. In our cryo-EM structure with 17 nt 5′-triphosphate RNA, RNA protrudes from the RNA exit channel and reaches the TPase of RNGTT (Figs. 1b and 2d), which belongs to the first branch of cysteine phosphatase. Docking with crystal structure of mouse RNA triphosphatase into TPase map ^40^,, the 5′ -triphosphate points towards the P loop of TPase, harboring a typical HCXXXXXRT motif and catalytic residue Cys126 in metazoan capping enzymes (Fig. 2d).

In the structure, the density of 15 nt RNA could be traced in the RNA channel, and the position of the 3^rd^ transcribed nucleotide is defined at RNA exit tunnel. Although there is extra density stretching to the bottom of TPase domain, the density is not clear enough to be identified. Its boundary is approximately 11.5 Å distance from the α-phosphate of the 3^rd^ transcribed ribonucleotide. Considering the 5′-triphosphate group, two additional ribonucleotides preceding the 3^rd^ ribonucleotide could theoretically be 18.5 Å, still not reaching Cys126 5 Å away. Nevertheless, interdomain movement of RNGTT and the flexibility of RNA are helpful for the enzymatic modules of RNGTT to move and catch the 17 nt RNA substrate, and complete enzymatic reaction (Fig. 2f, 2g).

Since γ-phosphates of 19 nt and 20 nt RNA extend from the 3^rd^ nucleotide to 30.5 Å and 36.5 Å, respectively (Fig. 2f), in a closer proximity to Cys126, RNGTT efficiently catalyzed the guanylation reaction (Fig. 2g). The 5′-triphosphate of 22 nt RNA is about 48.5 Å away from the 3^rd^ transcribed nucleotide and difficult to be accessed by the TPase of RNGTT (Fig. 2f) in consistent with the guanylation assay results (Fig. 2g). In spite of the long distance from the 5′ end of RNAs to the K294 active site of GTase, we speculated that the flexible linkage between two domains and the interdomain movement of GTase and OB fold would overcome the inaccessibility of the 5′-diphosphate of 17 nt, 19 nt and 20 nt RNA to K294 and facilitate accomplishing guanylation.

### Non-canonical positioning of NELF in PEC-RNGTT

In PEC-RNGTT cryo-EM data, we could separate RNGTT containing particles with NELF complex density (Supplementary Fig. 2b). NELF displays two different conformations (Fig. 3a, b). One conformation is similar to the previously reported PEC structure (canonical, “Up” state) (Fig. 3b), the other is downwardly positioned near the DNA duplex (non-canonical, “Down” state) (Fig. 3a). NELF-B-A-C/D models derived from PEC (PDB ID: 6GML) and NLEF-E from AlphaFold2 were rigidly fitted in cryo-EM maps (Supplementary Figs. 3e, f) and manually adjusted in Coot.

**Fig. 3.**
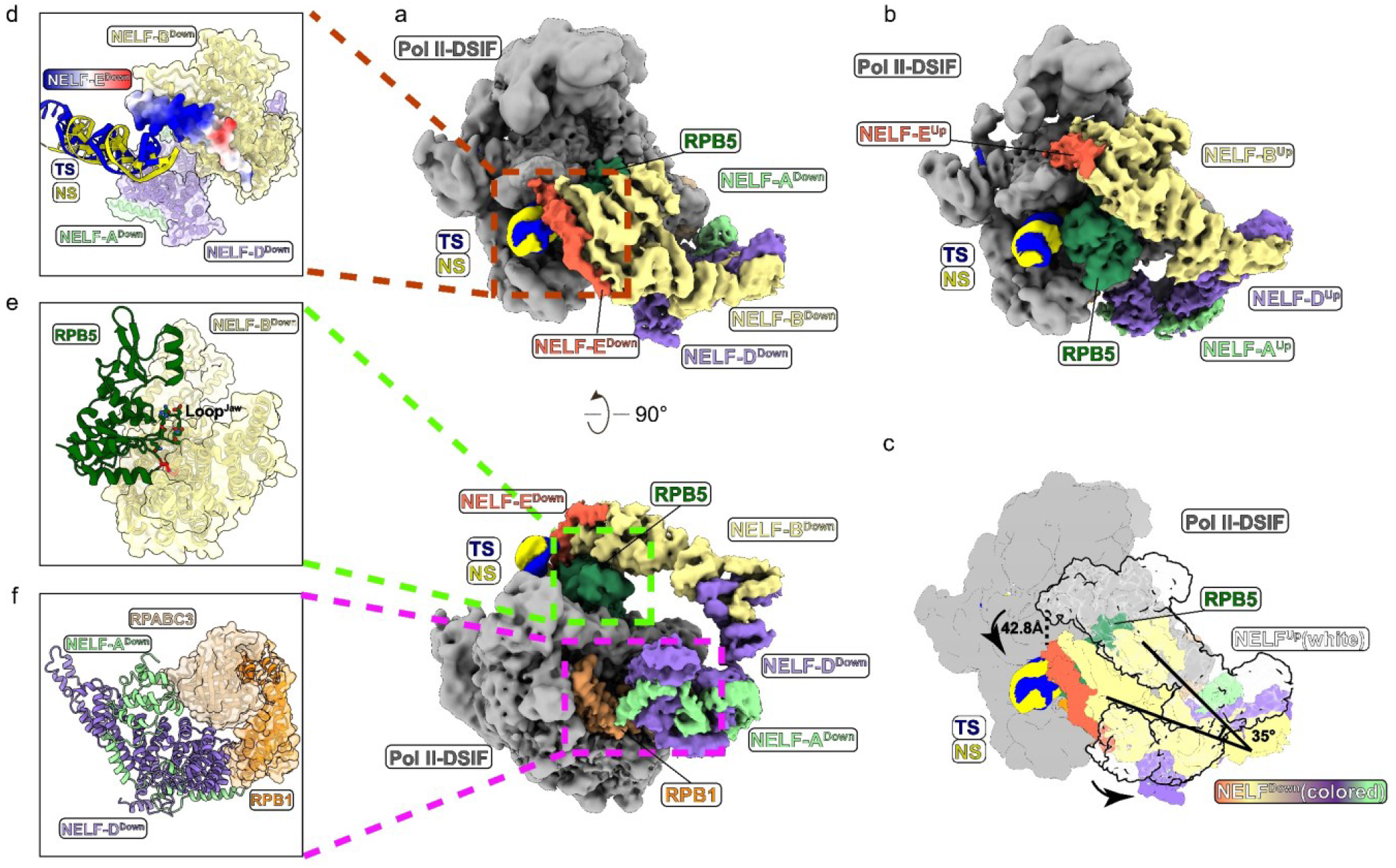
Structure comparison of NELF in two conformations. **a** Overall structure of NELF in “Down” state conformation in PEC-RNGTT in two views. The labels of different components are in corresponding map colors. **b** Overall structure of NELF in “Up” state conformation in PEC-RNGTT in the same view with **a** (above) corresponding labels around **c** Comparison of the cryo-EM maps of NELF “Down” state (colored) and “Up” state (white and transparent). Conformational difference is indicated by moving distance and angle arrows. The distance is measured referring to NELF-E N-terminal helix and the angle is measured by NELB central axis. **d-f** Close-up views of NELF^Down^ position on the Pol II. **d** N-terminal helix of NELF-E^Down^ is in close proximity to DNA. N-terminal helix of NELF-E^Down^ is displayed as a charged surface. **e** NELF-B^Down^ encloses RPB5 of Pol II. **f** Interaction between NELF-A^down^ and NELF-D^Down^ with RPB1 and RPABC3 of Pol II.

Comparing to the conformation in the “Up” state, the “Down” state conformation shows the following differences. (i) NELF complex is displaced as far as 42.8 Å at NELF-E N-terminal helix and rotates toward the entry DNA by 35 degrees (Fig. 3c). (ii) A previously undefined N-terminal region of NELF-E displays a longer helical density in “Down” state map, which was well fitted with NELF-E (1-37 aa) Alphafold2 model. It is a long positively charged helix possessing ten-Lysine residues. The helix is positioned near the entry DNA (Fig. 3d and Supplementary Fig. 4a). (iii) Instead of dangling on the loop of jaw domain in RPB5, the HEAT domain of NELF-B wraps the lateral surface of it (Fig. 3e and Supplementary Fig. 4b). (IV) In “Up” state, NELF-A neighbors RPB8, and NELF-D interacts with the Pol II funnel and trigger loop, which impedes Pol II progressing and retains Pol II pausing (Supplementary Fig. 4c). Due to NELF-A-NELF-D rearrangement, the N- and C-terminus of NELF-A-NELF-D lobe descends, the middle helix bundle is lifted to interact with RPB8. The C-terminal region of NELF-D detaches from RPB1 funnel and trigger loop, leaving the funnel partially open and convenient for NTP delivery to the active site (Fig. 3f).

### RNGTT and CMTR1 are arrayed on PEC surface

Given CMTR1 interaction network, genome-wide distribution and its role in the maturation of the 5′ end of pre-mRNA, CMTR1 has also been verified to function at early stage in pre-mRNA processing ^36^. Gradient centrifugation analysis showed that CMTR1 in sub-stoichiometry co-migrated with phosphorylated Pol II with DSIF, NELF in the absence or presence RNGTT (Supplementary Figs. 1c, f). We preferentially prepared cryo-EM sample of PEC-RNGTT-CMTR1, collected 10,690 micrographs and determined the complex structure at nominal resolution 4 Å with the periphery resolution from 5.5 Å to 10 Å (Supplementary Fig. 5d). The two conformations of NELF subcomplex were still observed in 3D classification (Supplementary Fig. 5b). We rigidly docked the crystal structure of RFM domain (PDB ID:4N48) ^14^ and AlphaFold2 model of GTase-like domain into CMTR1 cryo-EM map (Fig. 4a and Supplementary Fig. 3h), and structural models for other components of the complex were placed as described above (Supplementary Fig. 3g).

**Fig. 4.**
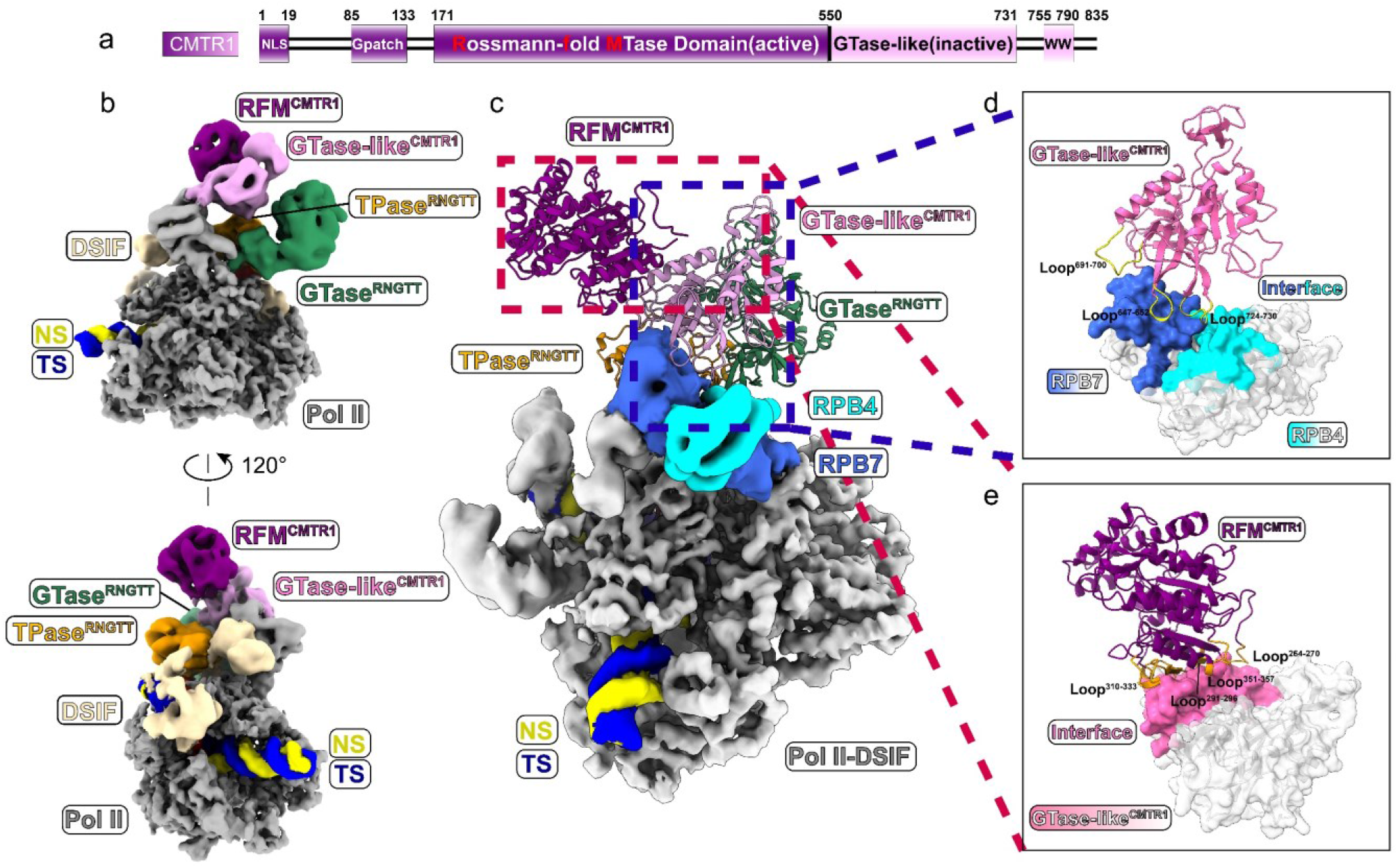
Cryo-EM structure of the PEC-RNGTT-CMTR1 complex. **a** Domain architecture of 2’-O-ribomethyltransferase, CMTR1. The cryo-EM map same color scheme is used throughout all figures. Solid black lines represent the linkage region between adjacent domains. **b** Combined cryo-EM map of PEC-RNGTT-CMTR1 complex. The labels of different components are in corresponding colors. **c** Ribbon models of RNGTT and CMTR1 dock on the PEC cryo-EM map. RFM and GTase-like domain of CMTR1 are displayed as models in dark magenta and plum respectively while other components are shown as density maps in corresponding tab colors. **d** Interaction between GTase-like domain of CMTR1 and Pol II stalk. Interface of RPB7 is highlighted as royal blue and interface of RPB4 is highlighted as cyan, while the other parts are white and transparent. Loops of GTase-like domain mentioned in the article are depicted as ribbons in yellow. **e** Interaction between RFM and GTase-like domain of CMTR1. Interface of GTase-like domain is highlighted as plum while the other part is white and transparent. Loops of RFM domain mentioned in the article are shown as orange ribbons.

In the PEC-RNGTT-CMTR1 structure, CMTR1 is anchored in the apical groove at the joint of RPB4 and PRB7 through flexible interaction, mediated by apical loop regions of RPB4 and RPB7. Three loops (647-652 aa, 691-700 aa, 724-730 aa) from GTase-like domain of CMTR1 contribute to CMTR1 docking on the Pol II tip (Fig. 4d). The connection between RFM and GTase-like domain was not restricted to the loop linkage between two domains, additionally, there is an intra-binding interface generated by GTase-like domain and RFM domain anchoring loops (264-270 aa, 291-296 aa, 310-333 aa, 351-357 aa) (Fig. 4e).

Comparison with the recently deposited structures of Pol II-DSIF-RNGTT-CMTR1 and Pol II-DSIF-CMTR1 (PDB ID:8P4E and 8P4F) ^45^ suggests that CMTR1 remains binding stalk through the GTase-like domain, when the RFM domain rotates to capture and methylate the substrate. In our PEC-RNGTT-CMTR1 structure, the full-length RNGTT was entirely observed, and TPase is still situated adjacent to the stalk (Fig. 5a). In contrast, TPase of RNGTT in one deposited structure (PDB ID:8P4E) ^45^ is invisible, and the entire RNGTT density is missing in the other one (PDB ID:8P4F) (Fig. 5b, c). Superimposed the three structures, the significant deviation derives from the angle of RFM domain in CMTR1, showing a clockwise rotation around the Pol II stalk (Fig. 5d). We speculated that the CMTR1 could dock onto Pol II in absence of RNA substrate in advance, then swing to search for the substate, catch the semi-capped pre-mRNA substrate and methylate it in the end. During this process, the two domains of RNGTT gradually clash with RFM domain in CMTR1, and turn out to be invisible.

**Fig. 5.**
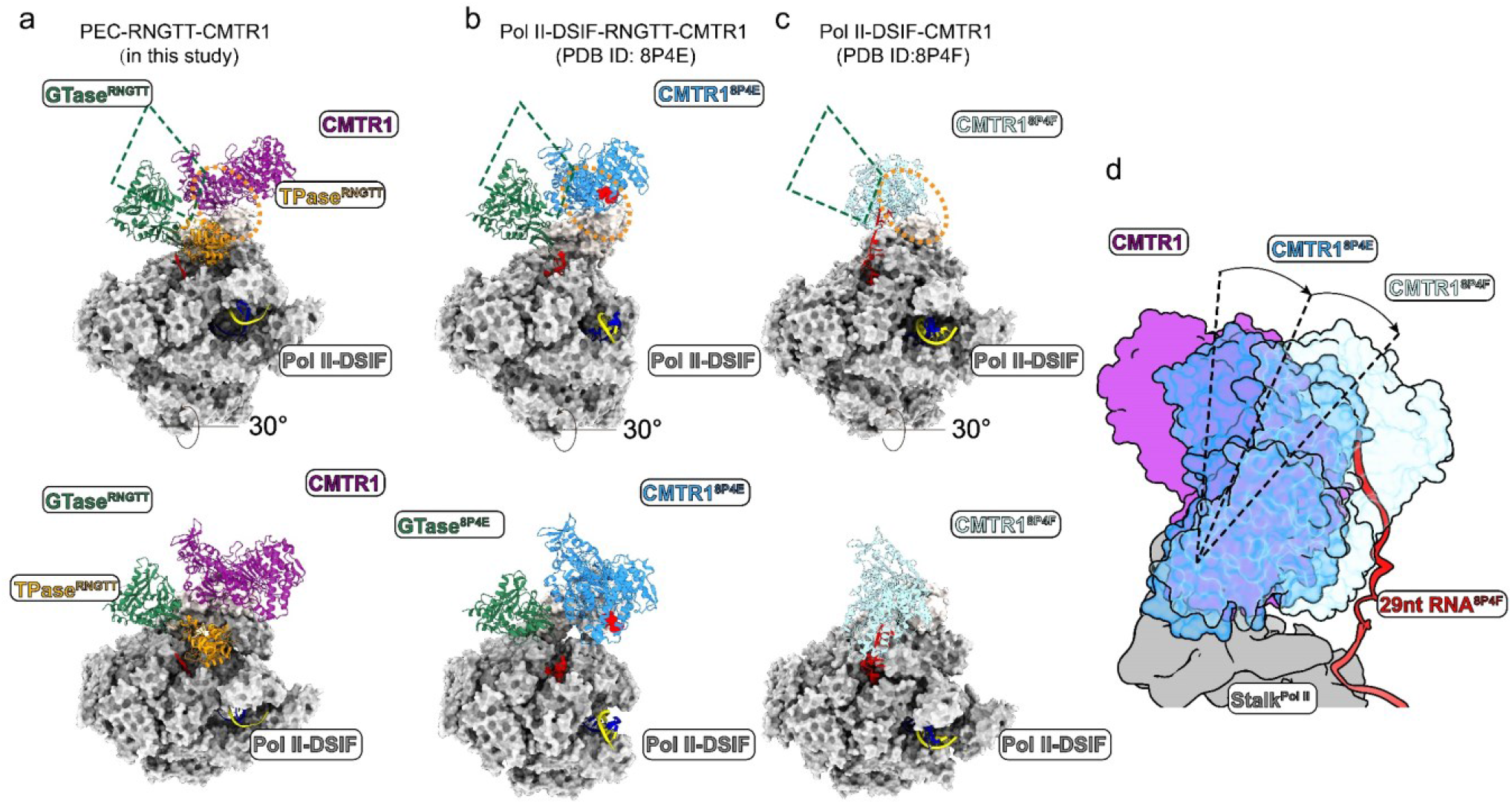
Structure comparison of PEC-RNGTT-CMTR1, transcribing Pol II-DSIF-RNGTT-CMTR1 and transcribing Pol II-DSIF -CMTR1. **a-c** Location of TPase^RNGTT^, GTase^RNGTT^ and CMTR1 of the three structures in two views. **a** PEC-RNGTT-CMTR1; **b** transcribing Pol II-DSIF-RNGTT-CMTR1 complex (PDB ID: 8P4E); **c** transcribing Pol II-DSIF-CMTR1 complex (PDB ID: 8P4F). **d** Orientation changes of CMTR1 in PEC-RNGTT-CMTR1 complex, 8P4E and 8P4F. CMTR1 in PEC-RNGTT-CMTR1 complex is colored in solid magenta, while CMTR1 in 8P4E and 8P4F in transparent sky blue and light blue, respectively.

## Discussion

In this study, we determined the structures of PEC-RNGTT and PEC-RNGTT-CMTR1. In both datasets, 3D classifications dissected Pol II bound with RNGTT and NELF complex. RNGTT and NELF complex are located at different positions of Pol II, RNGTT adjacent to the stalk and NELF around the foot and funnel domains of RPB1. There is no observable direct interaction between them in our structures.

Interaction between RNGTT and phosphorylated Ser5-CTD was not captured in our structure. Instead, we observed OB fold of RNGTT inserting into the cleft at root of the RPB4-RPB7 stalk. The GTase domain preceding the OB fold drifts outwards. This configuration indicates that mobility of GTase, rather than OB fold, accounts for the open/closed conformation transition of cleft, for OB fold is tightly clamped by stalk and RPB1. The movement of GTase also contributes to catch the 5′-diphosphate RNA for guanylation. We speculated the interaction of OB fold with Pol II is strong enough to maintain RNGTT localization when RNGTT adopts different conformations to exert its dephosphorylation and guanylation activity.

The interaction between N-terminal region of TPase and OB fold bent the RNGTT into a circular configuration with an interspace probably for shift of GTase and accommodation of the released 5′-diphosphate RNA. Moreover, this interaction facilitates the TPase catalytic pocket facing towards the RNA exit tunnel. Compared with the PEC structure ^43^, TPase substitutes for the position of KOWx-KOW4 in our structure, whose density is not observed in our cryo-EM maps, presumably because the KOWx-KOW4 is jostled to another place or changes its conformation. At this position, TPase is able to access the 5′-triphosphate of the nascent pre-mRNA.

The minimal RNA length of capping substrate varies in different capping assays. *In vivo* assays using nuclear extract have demonstrated substrate length covers a wide range of 20 nt-79 nt ^22^, and up to 223 nt RNA could be capped *in vitro* recombinant ternary complex assay ^28^. We set up a recombinant guanylation assay *in vitro*, showing that the length of 17 nt is enough to be processed and consistent with previous result ^20^. However, it is not the exclusive length, as long as P loop of TPase is capable to catch the substrate, it could catalyze the dephosphorylation reaction. 17 nt, 19 nt and 20 nt pre-mRNA could be processed, but not 22 nt. It is perhaps regulated by the moving range of nascent RNA, and the mobility of TPase influenced by TPase-GTase linker flexibility and its binding affinity to the OB fold. We used four *in vitro* transcribed RNA hybridized with template DNA as the scaffold to assemble the Pol II elongation complex in the presence of DSIF, without further purification of the artificial complex, which possibly leads to inefficient capping of 22 nt RNA.

In both of PEC-RNGTT and PEC-RNGTT-CMTR1 cryo-EM maps, we observed two conformations of NELF subcomplex. Compared with the canonical conformation in PEC ^43^, the new conformation exhibits obvious distinction from the canonical one, particularly in NELF-A-NELF-C/D lobe. The rearrangement of NELF-A and NELF-D helices dissociate NELF-D from Pol II funnel and trigger loop, half opening the gate for NTP entry.

CMTR1 is crucial for 5′ end processing prior to nuclear export. It firmly anchors at the tip of Pol II through the loop regions of GTase-like domain, enabling the RFM domain to swing around and search for its substrate. Furthermore, GTase-like domain also intra-molecularly interact with RFM domain. Considering its effect on CMTR1′ s enzymatic activity ^14^, it is likely that GTase-like domain could stabilize the RFM domain to be active. As RNA elongating, the entire CMTR1 molecule rotates around the anchoring point of the stalk, potentially aiding the assembly of other transcription factors onto the Pol II transcription machinery.

Till now, structures of eukaryotic co-transcriptional capping enzyme have been reported in two papers^39,45^. One of the structures is the *S. cerevisiae* phosphorylated Pol II with CE void of transcription factors, which was determined at a nominal resolution 17.4 Å (EMD-2922) ^39^. By means of native mass spectrometry, hetero-trimeric and hetero-tetrameric CE are identified on the Pol II surface, and both CEs could dephospharylate and guanylate the 23 nt RNA substrate. Although the resolution is limited, the map shows that CE spans the RNA exit tunnel and OB domain of Ceg1 forms a small contact with the Rpb4/7 subcomplex.

Compared with the lately reported human co-transcriptional capping complex^45^, we coincidentally assembled the PEC-RNGTT and PEC-RNGTT-CMTR1 complex with the same components, including the transcription factors. For the PEC-RNGTT, we employed 17 nt RNA to determine its structure and observed the full-length RNGTT anchoring in close proximity to the Pol II stalk, also reported in the latest paper, but not released in databank yet. Furthermore, we detected two conformations of NELF complex in the PEC-RNGTT structures. The prominent rearrangements of NELF-A-NELF-D lobe in the “Down” state result in dissociation of NELF complex from Pol II funnel and trigger loop, implying the transitional situation of Pol II from pausing to release or some other events, for example late pausing ^19^ and cap binding ^46^. For the PEC-RNGTT-CMTR1, we captured both full-length RNGTT and CMTR1 visibly arrayed adjacent to the Pol II stalk. The catalytic domain of CMTR1 is further away from the RNA exit tunnel than latest two structures of transcribing Pol II with CMTR1(PDB ID:8P4E and 8P4F) ^45^, probably because the longer RNA we used is difficult for CMTR1 to catch and render us not to find the RNA density, implying it is possibly an initial position for CMTR1 staying on the Pol II prior to substrate recognition.

Unfortunately, the methyltransferase RNMT-RAM failed to be assembled into PEC or any other form of Pol II. The mechanism of RNMT-RAM methylation remains elusive. In consideration of the dependence of RNA during RNMT-RAM functions, it may also require a just suitable RNA in certain Pol II stage. In our PEC-RNGTT-CMTR structure, there is enough space to accommodate extra proteins. Nevertheless, it′s not excluded that some unknown factors will help. Secondly, NELF-E plays important roles in NELF recruitment, cap-binding of CBC and 3′ end processing^46-48^. The essential NELF-E tentacle is so flexible that we only observed its N-terminal helix. The limited structural information hampers the understanding of NELF-E functions during the co-transcriptional capping process. Taken together, our cryo-EM structures shed light on the co-transcriptional pre-mRNA capping and methylation to some extent, particularly correlate capping with Pol II paused elongation complex and captured several conformational changes of enzymes and transcription factors.

## Methods

### Protein expression and purification

Pol II was purified from *S. scrofa* thymus and transcription factor DSIF and NELF were prepared as previously described ^31^. There are four residue substitutions (G882S of RBP2, T75I of RPB3, S140N of RPB3, and S126T of RPB6) between *S. scrofa* and *H. sapiens* Pol II. Full-length genes of human RNGTT and CMTR1 were cloned into pMlink vector containing a 4×ProteinA tag at the N-terminal and were transfected to Expi293F cells (Thermo Fisher Scientific) when cell density is 2.5× 10^6^ /ml. Cells were harvested by centrifugation after cultured for another 72 h at 37 °C. Protein purification were performed at 4°C. Cells were lysed for 30 mins in buffer A (30 mM HEPES-NaOH, pH 8.0, 300 mM NaCl, 10% (v/v) Glycerol, 0.25% (w/v) CHAPS, 5 mM MgCl_2_, 5 mM ATP, 2 mM DTT (Dithiothreitol), 1 μg/mL Aprotinin, 1 μg/mL Pepstatin, 1 μg/mL Leupeptin. Cell lysate was clarified by centrifugation for 30 mins at 16,000 rpm using JLA-16.250 rotor (Beckman Coulter). The supernatant was kept and incubated with equilibrated IgG resin (Smart-Lifesciences) for 6 hours and washed thoroughly with buffer B (30 mM HEPES-NaOH, pH 8.0, 300 mM KCl, 10% (v/v) Glycerol, 0.1% CHAPS, 2 mM MgCl_2_, 5 mM ATP, 2 mM DTT). Target protein was released from the resin after on-column digestion overnight and eluted with buffer B. RNGTT was further loaded onto Mono Q (5/50GL, GE Healthcare) while CMTR1 was further loaded onto a Heparin column (5/50GL, GE Healthcare). Both proteins were eluted from the columns with NaCl concentration gradient. The peak fractions were collected and concentrated to ∼1 mg/mL respectively. The concentrated samples were preserved at –80°C for subsequent biochemical and structural analyses.

### Pol II phosphorylation

Ser5-phosphorylated Pol II was catalyzed by transcription factor TFIIH, which was prepared as described previously ^49^ and immobilized on IgG resin where the phosphorylation reacted. The TFIIH-contained resin was incubated with Pol II at 25°C for 15 mins in buffer C (30 mM HEPES-NaOH, pH 8.0, 100 mM KCl, 5% (v/v) Glycerol, 6 mM MgCl_2_, 50 μM ATP, 2 mM DTT for final concentration. Phosphorylated Pol II was eluted with buffer E and quantitated using SDS-PAGE gel stained by Coomassie blue. The protein was stored at –80°C for further experiments. All the Pol II mentioned below was phosphorylated unless specially noted.

### Preparation of 5′-triphosphate oligoribonucleotides

5′-triphosphate oligoribonucleotides was synthesized by *in vitro* transcription using T7 High Yield RNA Synthesis Kit (YEASEN). Transcription template and HDV ribozyme with T7 promoter sequences were cloned into pUC57 vector and the extracted plasmids were further purified with Source Q 5/50 GL column (GE Healthcare). The DNA fractions were precipitated with 0.3 M NaAc and 70% (v/v) isopropanol and recovered (Sangon Biotech). The plasmids were linearized using XhoI (NEB) and extracted by phenol/chloroform (1:1). The final concentrations of the templates were measured about 1 μg/μl. All the steps below were RNase-free. According to the specification of the kit, *in vitro* transcription was performed. The transcription reaction was incubated at 60°C for 30 mins to release target oligoribonucleotides from HDV ribozyme. The products were loaded onto 20% denaturing gel (7 M urea, 1×TBE, 20% Bis-tris acrylamide 19:1 gel). The target RNA bands were excised from gels and soaked in 0.3M sodium acetate overnight at 4°C. The RNA was precipitated with isopropanol. The precipitates were dissolved in DPEC water (Sangon Biotech) and analyzed with 15% native PAGE (1×TG, 15% Bis-tris acrylamide 19:1 gel) gel and nanodrop.

### Sample preparation for cryo-EM

To obtain the DNA−RNA hybrid, template strand DNA (Generay Biotechnology) and 5′-triphosphate RNA were mixed with a molar ratio of 1:1.3 and were annealed following 95 °C for 5 mins and then decreasing the temperature by 1°C min^−1^ steps to 4°C in a thermocycler in 20 mM HEPES-KOH, pH 7.4, 60 mM KCl, 3 mM MgCl_2_, and 5% (v/v) glycerol. All concentrations below referred to the final concentrations used in the complex assembling. Phosphorylated Pol II (120 pmol) was incubated with DNA-RNA hybrid at a 1:1.2 molar ratio for 10 mins at 30°C. Hybrid containing17 nt 5′-triphosphate RNA was used for PEC-RNGTT samples, and hybrid containing 40 nt 5′-triphosphate RNA was used for PEC-RNGTT-CMTR1 sample. Then non-template DNA was added at a 1:1.2 molar ratio with hybrid followed by another 10 mins incubation at 30°C, producing an elongation complex (EC). Further assembly was stepwise and carried out at 25°C for 20 mins. Four-fold molar excess of DSIF (480 pmol) was added firstly, and four-fold molar excess of capping enzymes (480 pmol), RNGTT alone or with CMTR1 were complemented subsequently. Then four-fold molar excess of NELF (480 pmol) was added.

The assembled complexes were purified and crosslinked using gradient fixation (Grafix) ^50^. The homogeneity of peak fractions was assessed by negative-staining electron microscopy. Qualified fractions were pooled, concentrated and replaced buffer to reduce the glycerol concentration under 0.5% (v/v). Negative-staining EM grids were prepared as previously described ^51^. For cryo-EM grids preparation, 3 μL of the concentrated samples were applied to Quantifoil R 1.2/1.3 holey, 200 mesh carbon grids which are freshly glow-discharged in the H_2_/O_2_ mixture for 30 s using a Gatan 950 Solarus plasma cleaning system with a power of 5 W. After incubation of 5 s at a temperature of 4°C and a humidity of 100 %, the grids were blotted for 2 s with blot force -2 in a Thermo Fisher Scientific Vitrobot Mark IV and plunge-frozen in liquid ethane at liquid nitrogen temperature. The ø55/20mm blotting paper (TED PELLA) is used for blotting.

### *In vitro* capping assay

*In vitro* capping assay was performed using purified RNGTT and EC containing 5′-triphosphate RNA as substrate. EC with different length RNAs (17 nt, 19 nt, 20 nt, 22 nt) was assembled as above, and the molar ratio between phosphorylated Pol II and hybrid was adjusted to 1.2:1. For one reaction, EC (3 pmol defined by hybrid) was mixed with RNGTT (8 pmol) in buffer containing HEPES-NaOH, pH 8.0, 100 mM NaCl, 5 mM MgCl_2_, 2 mM DTT. The reactions were started with the addition of 100 μM rGTP at 37°C for 60 mins and stopped by adding the same volume of stop buffer containing 7 M urea, 50 mM EDTA and 1×TBE. 1μl of Protease K (Promega, 20 mg/ml) was added and digested for 30 mins at 30°C. The reaction samples were analyzed by 20% denaturing gel (8 M urea, 1×TBE, 20% Bis-Tris acrylamide 19:1 gel) in 1x TBE buffer for 50 mins at 180 V constant. The gels were stained with Sybr™ Gold dye (Invitrogen, ThermoFisher Scientific) and visualized using Typhoon 9500 FLA Imager (GE Healthcare Life Sciences).

### Cryo-EM data collection

The cryo-EM data collection was finished with a Thermo Fisher Scientific Titan Krios transmission electron microscope operated at 300 kV. Cryo-EM images were automatically recorded by a Gatan K3 Summit direct electron detector equipped with a GIF quantum energy filter (Gatan) set to a slit width of 20 eV. All images were collected in the super-resolution counting mode using Serial-EM with a nominal magnification of ×64,000 in the EFTEM mode, yielding a super-resolution pixel size of 0.667 Å on the image plane. The defocus values ranging from −1.5 to −2.5 μm. Each micrograph stack was dose-fractionated to 40 frames with a total electron dose of ∼50 e−/Å2 and a total exposure time of 3.6 s. 15281 micrographs of phosphorylated PEC-RNGTT and 10690 micrographs of phosphorylated PEC-RNGTT-CMTR1 were collected for further processing.

### Image processing

Movie stacks were aligned by MotionCor2 ^52^ with 5×5 patches and 2x binned to a calibrated pixel size of 1.334 Å/pixel, generating drift-corrected micrographs with and without electron-dose weight. Contrast transfer function (CTF) parameters were estimated by Gctf ^53^ from non-dose weighted micrographs. The particles were automatically picked by Gautomatch. The data processing were performed with RELION3.1^54,55^ and cryoSPARC ^56^ v4 using dose-weighted images.

For PEC-RNGTT dataset, the autopicked particles were extracted with 320^3^ pixel box size and rescaled the particles to 80^3^ pixel box size. The whole particle dataset was split into several sub-datasets for further reference-free 2D classification. The yielded particles were further subjected to the 3D classifications. The particles with good quality were rescaled to 160^3^ pixel box size and subsequently subjected to the 3D classification. This process provided two classes with different NELF conformations. The particles from the two NELF conformations were subjected for further 3D classification with different NELF module masks. A final set of 51,449 and 68,862 particles for NELF in “Up” state and NELF in “Down” state were selected to perform a final 3D reconstruction in cryoSPARC, yielding reconstruction of overall maps with “Up” state NELF and “Down” state NELF at. Local refinement focused on NELF module with mask could reconstituted the NELF part at 4.62 Å and 4.83 Å, respectively. To improve the map quality of the RNGTT, the signal of RNGTT and partial of the Pol II stalk module were subtracted from six classes of 3D overall classification. The subtracted particles were subjected for further 3D classification by applying local mask for RNGTT and stalk module. The yielded particles were further subjected for particles subtraction with RNGTT mask. A 3D classification by applying mask of the RNGTT resulted in a clean dataset containing 175,576 particles. The resulting particles were refined in cryoSPARC, yielding a reconstruction of RNGTT at 5.82 Å.

PEC-RNGTT-CMTR1 dataset were processed as described above.

### Model building and structure refinement

Model building was carried out by fitting the available cryo-EM and crystal structures of Pol II, NELF and RNGTT from PDB: 6GML, TPase of RNGTT from PDB: 1I9S, GTase PDB:3S24 and RFM domain of CMTR1 from PDB: 4N48 into the EM density maps using UCSF Chimera. The models of GTase-like domain of CMTR1 and the long helix of NELF-E were predicted by AlphaFold2^57^. The model was then manually adjusted in Coot ^58^. The final model refinement was carried out using phenix.real_space_refine with PHENIX ^59^ and validated through examination of Ramachandran plot statistics, a MolProbity score. Model representations in the figures were prepared by PyMOL (http://pymol.org/) and UCSF ChimeraX ^60^.

## Acknowledgement

We thank Dr. Xizhi Chen, Jiabei Li and Xinxin Wang for preparing Pol II from *S. scrofa* thymus. We thank Dr. Hai Zheng for RNA-DNA hybrid preparation. We thank the Center of Cryo-Electron Microscopy at Fudan University for the supports on cryo-EM data collection. This work was supported by grants from the National Natural Science Foundation of China (31630002) and Shanghai Pujiang Program (21PJD004).

## Author Contributions

Y.L. and Z.L. prepared Pol II in the apo form for structural analyses; Y.L. purified the other proteins with the help from Z.L.; Z.L. designed and prepared nucleic acid scaffold; Y.L. assembled the complexes, prepared cryo-EM samples and collected the data; Z.L. and Q.W. performed the cryo-EM analyses and the model buildings; Y.L. and Z.L. performed the capping assays and analyzed the data; Y.X., Z.L. and Y.L. designed the experiments and analyzed the data; Z.L.,Y.L. and Q.W. wrote the manuscript; and Z.L. supervised the project.

