## Supplementary Material for "Structures of co-transcriptional RNA capping enzymes on paused transcription complex"

### Supplementary Figures

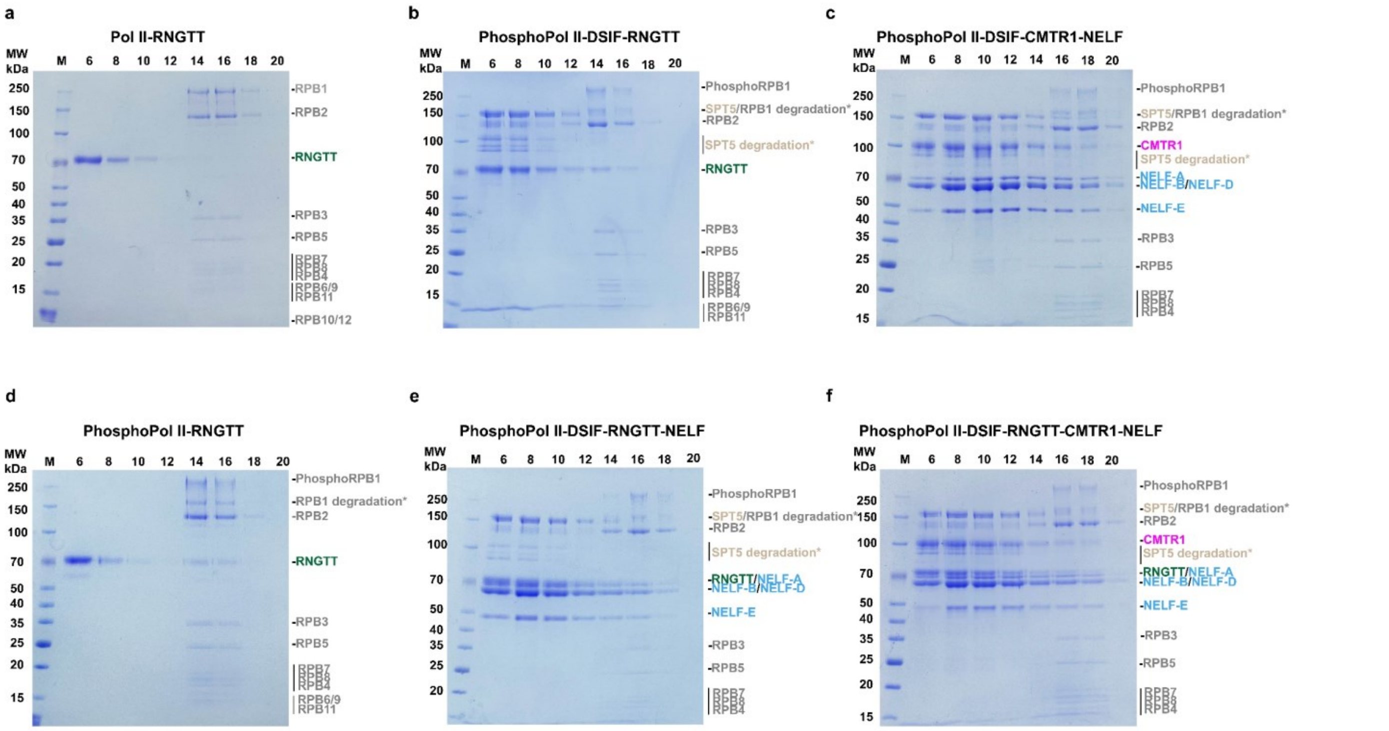


**Fig. S1 Complex assembly through glycerol gradient ultracentrifugation. a-f** Indicated fractions of the assembly of complexes containing different components through glycerol density gradient ultracentrifugation.

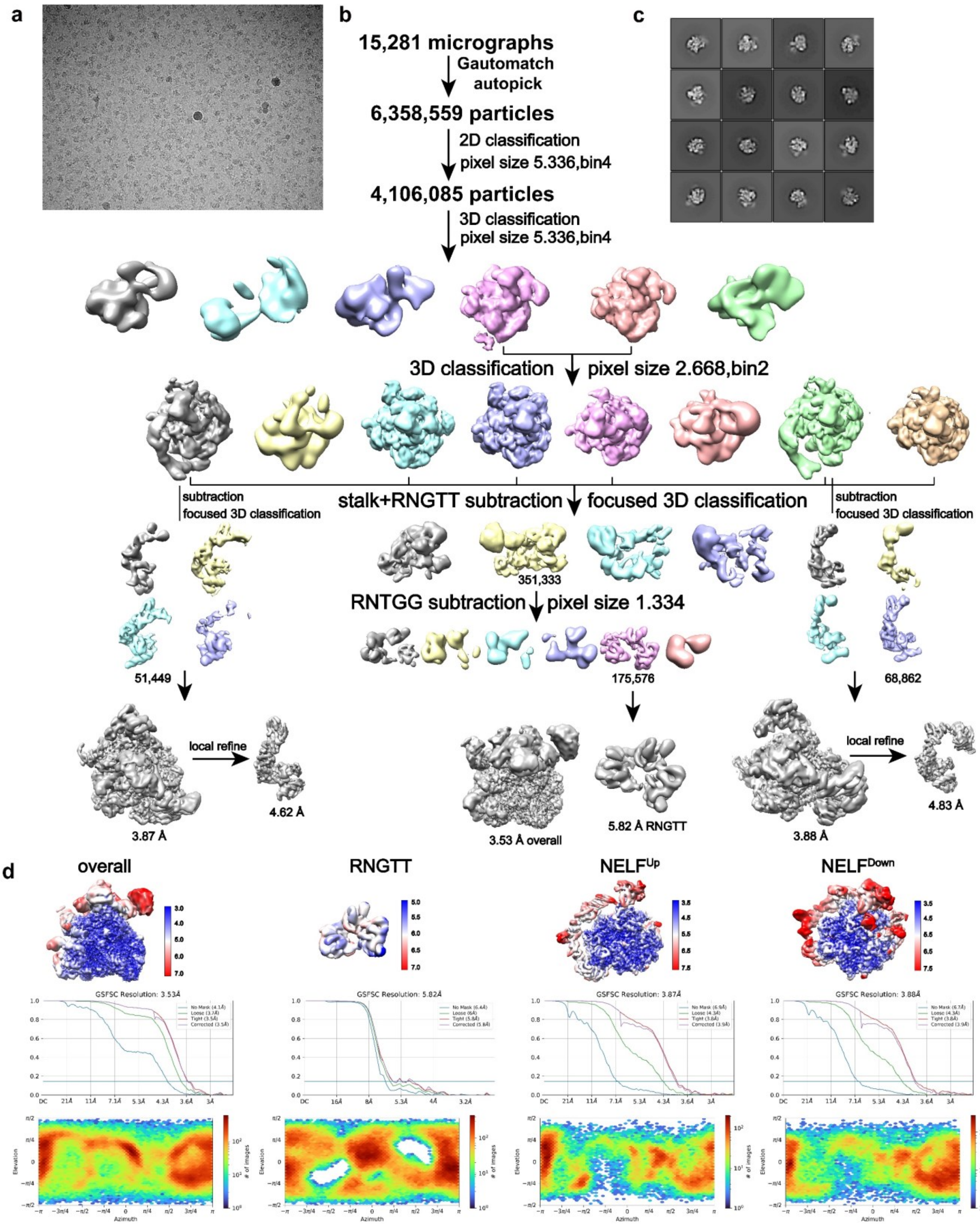


**Fig. S2 Data collection and image processing of PEC-RNGTT complex. a** Representative cryo-EM micrograph. **b** Flow-charts of the cryo-EM image processing and 3D reconstructions of the human PEC-RNGTT. **c** 2D classification from at least three times repeatedly of the human PEC-bound RNGTT. **d** Local resolution estimation, FSC curves and orientation of the cryo-EM reconstructions of Pol II-DSIF-RNGTT complex, masked RNGTT, PEC-RNGTT complex with NELF$^{Up}$ conformation and PEC-RNGTT complex with NELF$^{Down}$ conformation.

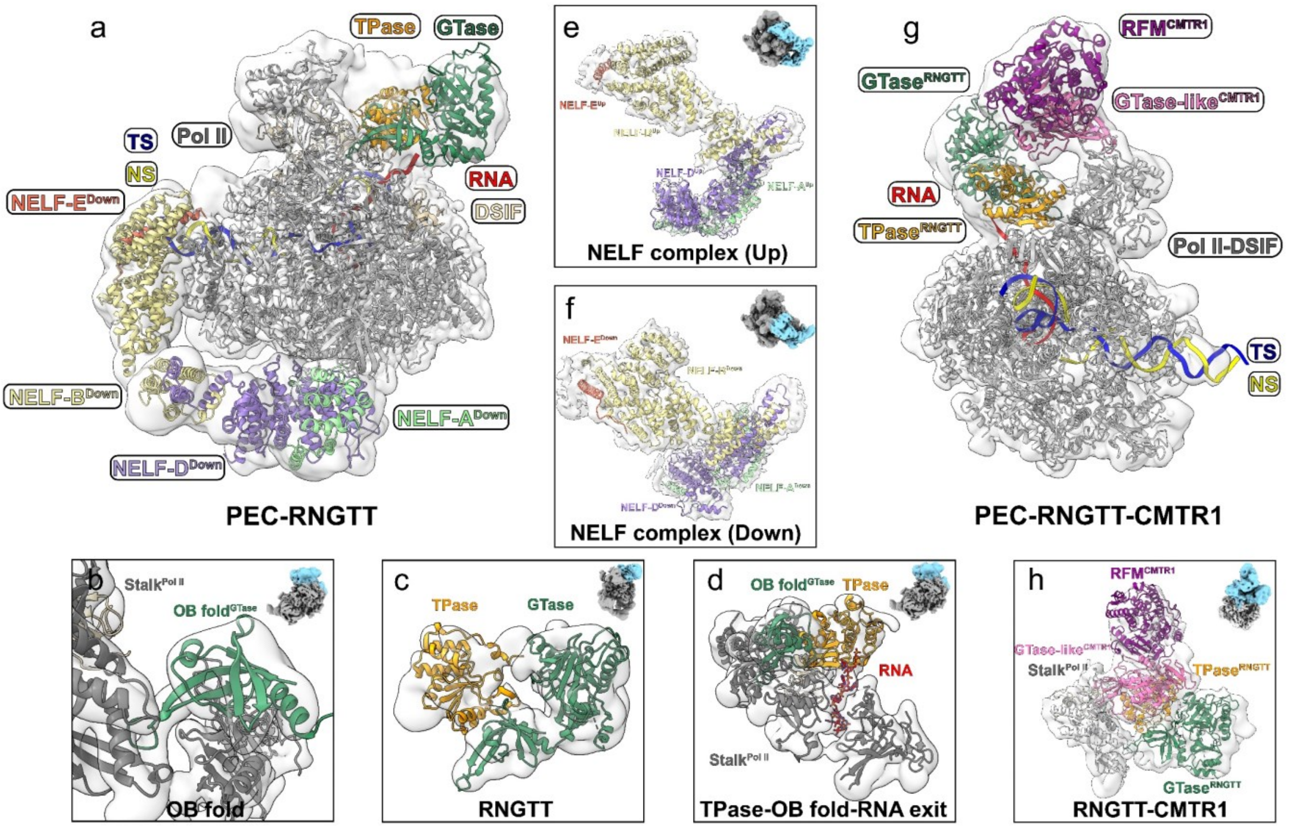


**Fig. S3 Cryo-EM maps and structural models of PEC-RNGTT and PEC-RNGTT-CMTR1 complex. a, g** Overall cryo-EM maps of PEC-RNGTT(**a**) and PEC-RNGTT-CMTR1 complex(**g**) fitted with individual model. Cryo-EM maps are shown in transparent surfaces with models shown in cartoon. **b-f, h** Focused refinement cryo-EM maps in different regions are shown in boxed panels. The maps at the top right indicate the masked regions. The structural models fit into the maps well, indicating correct placement of the structural models.

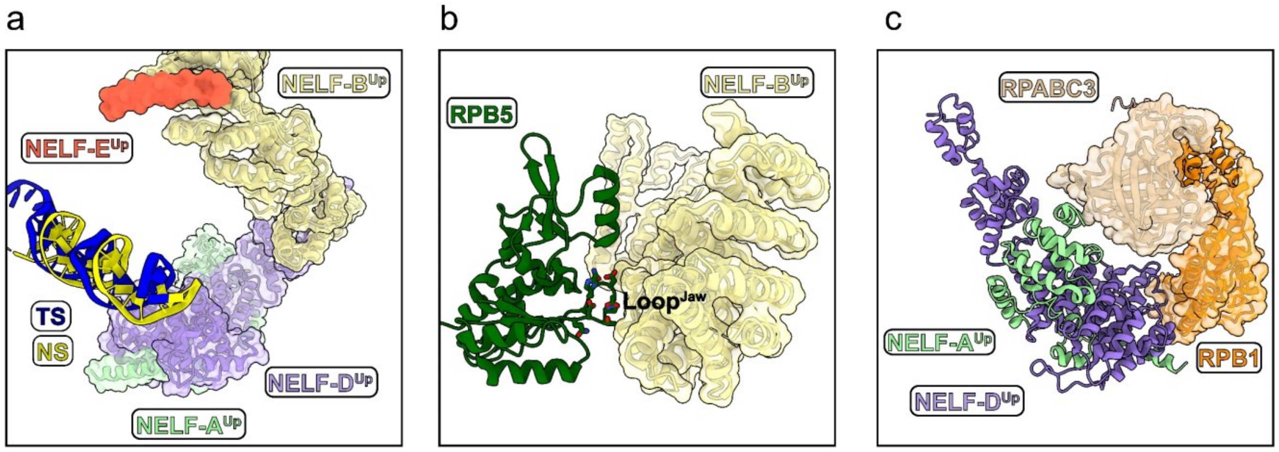


**Fig. S4 Close-up views of NELF and Pol II interactions in the "Up" state conformation. a** Architecture of NELF-E$^{Up}$ subunit and its positional relationship with DNA. **b** Interaction between NELF-B$^{Up}$ and RPB5 of Pol II. **c** Interaction between NELF-A$^{Up}$ and NELF-D$^{Up}$ with RPB1 and RPABC3 of Pol II.

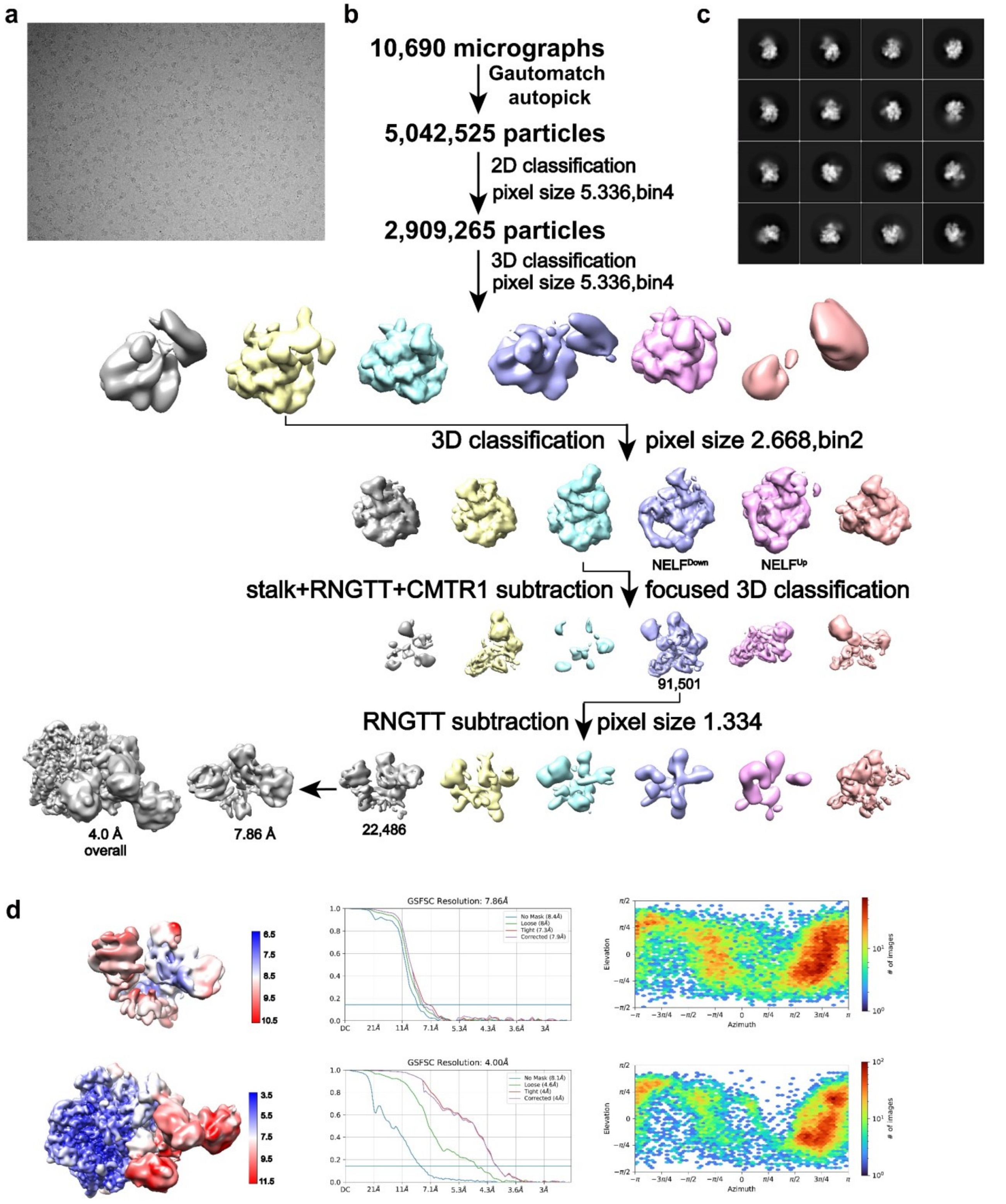


**Fig. S5 Data collection and image processing of PEC-RNGTT-CMTR1 complex. a** Representative cryo-EM images. **b** Flow-charts of the cryo-EM image processing and 3D reconstructions of the human PEC-RNGTT-CMTR1 complex. **c** 2D classification from at least three times repeatedly of the human PEC-RNGTT-CMTR1 complex. **d** Local resolution estimation, FSC curves and orientations of the cryo-EM reconstructions of PEC-RNGTT-CMTR1 and focused refinement RNGTT-CMTR1.
